# The statistical power of three monkeys

**DOI:** 10.1101/2022.05.10.491373

**Authors:** Jean Laurens

**Affiliations:** Ernst Strüngmann Institute (ESI) for Neuroscience, Frankfurt, Germany

## Abstract

Neuroscience studies in non-human primates (NHP) often follow the rule of thumb that results observed in one animal must be replicated in at least one other. However, we lack a statistical justification for this rule of thumb, or an analysis of whether including three or more animals is better than including two. Yet, a formal statistical framework for experiments with few subjects would be crucial for experimental design, ethical justification, and data analysis. Also, including three or four animals in a study creates the possibility that the results observed in one animal will differ from those observed in the others: we need a statistically justified rule to resolve such situations. Here, I present a statistical framework to address these issues. This framework assumes that conducting an experiment will produce a similar result for a large proportion of the population (termed ‘representative’), but will produce spurious results for a substantial proportion of animals (termed ‘outliers’); the fractions of ‘representative’ and ‘outliers’ animals being defined by a prior distribution. I propose a procedure in which experimenters collect results from M animals and accept results that are observed in at least N of them (‘N-out-of-M’ procedure). I show how to compute the risks α (of reaching an incorrect conclusion) and β (of failing to reach a conclusion) for any prior distribution, and as a function of N and M. Strikingly, I find that the N-out-of-M model leads to a similar conclusion across a wide range of prior distributions: recordings from two animals lowers the risk α and therefore ensures reliable result, but leaves a large risk β; and recordings from three animals and accepting results observed in two of them strikes an efficient balance between acceptable risks α and β. This framework gives a formal justification for the rule of thumb of using at least two animals in NHP studies, suggests that recording from three animals when possible markedly improves statistical power, provides a statistical solution for situations where results are not consistent between all animals, and may apply to other types of studies involving few animals.

## Introduction

When performing experiments on animals, choosing the proper number of animals involves a trade-off: on the one hand, ethical and practical considerations compel experimenters to reduce the number of animals. On the other hand, using too few animals endangers the validity and reproducibility of experimental results. Statistical methods offer a rigorous framework to guide this decision.

In the context of neuroscience experiments in non-human primates (NHPs), choosing the right number of animals involves specific challenges. Ethical and practical considerations often limit experimenters to a few animals, typically two or three. Using so few animals is possible because neuroscience studies in NHPs generally aim at discovering fundamental aspects of cognition or physiology that are expected to be evident in small numbers of animals (as opposed to measuring quantitative effects, which necessarily involve larger samples). Yet, even basic anatomy, physiology or behaviour can vary between animals, and therefore a certain amount of duplication is required to establish a result. To address this issue, neuroscientists working on NHPs have often adopted the rule of thumb that results observed in one animal should be reproduced in a second one.

Yet, we lack a statistical framework to justify and interpret this rule of thumb. What do we truly gain by using two animals instead of one? Would using three or four animals confer any additional advantage? These questions have fundamental implications for experiment planning, data interpretation and reviewing, and ethical discussions.

In here, I propose an answer to these questions by developing a framework called ‘N-out-of-M’ (N-oo-M) for interpreting and drawing inferences from experiments performed on small numbers (M) of animals. Within this framework, I define and calculate ‘α’ risks (defined here as the probability of reaching a false conclusion) and ‘β’ risks (defined here as the probability of failing to reach a conclusion), under a variety of scenarios. I find that using two animals generally yields reliable conclusions (i.e. low α) but will often lead to inconclusive results (i.e. high β), whereas using three animals can offer a sensible balance of α and β risks.

## Fundamental assumptions

### This framework is based on two principles

First, it applies to experiments that aim at establishing fundamental structural properties (e.g. the existence of place cells or mirror neurons in a cortical region, or whether an optogenetic stimulation affects motricity, or to delineate a cortical region) rather than quantitative measures (e.g. the average and standard deviation of Macaque’s body weight).

Second, this framework is based on the concept of ‘outlier’ animals. In this framework, ‘outliers’ refers to animals in which a certain result can be observed clearly and consistently, even though it is not observed in most of the population. For instance, an animal may develop an atypical behavioural strategy to solve a task, leading to the appearance of neuronal responses that can’t be reproduced in other animals. Or an animal may have anatomical anomalies or have experienced an undetected pathological event that affects an experiment’s outcome. Finally, technical errors (such as electrode misplacement) can affect the experiment outcome. In all cases, this may lead experimenter to observe incorrect results.

The potential for outliers leads to what I define as a risk α in this framework: the risk of reaching a conclusion that doesn’t generalize to the population, due to experimental variability. For instance, if 10% of animals have an abnormal behaviour which is not representative of the population as a whole, then recording from a single animal creates a risk α of 10%. Critically, the risk α accumulates when there is more than one possible type of outliers. If 5% of animals have abnormal behaviour, and 5% of animals have abnormal anatomies, and 5% of experiments are affected by errors, then the total risk of selecting an outlier animal is about 15%, which is unacceptably high.

This framework explores how using more than one animal may help lowering the risk α, i.e. ruling out outliers. For instance, one may record from two animals and only accept experimental outcomes that are observed in both. Since the probability that two animals are outliers is low, the risk α is reduced. But how much exactly is it? Furthermore, recording from multiple animals raises the question of what to do when the results observed in different animals don’t agree. For instance, could we design a strategy where we record from 3 animals and accept any outcome that is observed in at least two on them? What would be α risk associated with this strategy? I will show how to compute it, and discuss the results.

## Statistical model

Formally, this framework is defined by the following three hypotheses:

- Hypothesis 1: we assume that, in each animal, an experiment can lead to a number of *qualitatively* distinct outcomes (named A, B, C, etc). An ‘outcome’ may refer to a physiological, anatomical, behavioural result: it may be that a brain region contains a sizeable population of place cells, or that gamma oscillations in the visual cortex increase markedly when viewing a grating, or that a given brain nucleus projects onto another, or than a monkey learns to open a bottle to retrieve a treat. An outcome may also be a null result.
- Hypothesis 2: we assume a *prior* distribution across all possible outcomes. For instance, in a recording experiment aiming at the hippocampus, one may observe place cells in 80% of animals, grid cells 5% of animals (due to an error in electrode placement), another type of neurons in 5% of animals (due to atypical behaviours) and nothing in another 10%. In this situation, we call the most likely outcome the ‘representative’ outcome and other ‘outliers’ (we will also extend the model to cases where more than one outcome is ‘representative’).
- Hypothesis 3: I will now describe the ‘N out of M’ (N-oo-M) model. In this model, the experimenters perform the experiment in M animals. If they observe the same outcome in at least N animals, they will accept that it is the representative outcome. Otherwise, the experiment is inconclusive. This can lead to 3 results:

○ A correct conclusion, where the experimenters correctly identify the representative outcome.
○ An error, where one outlier is observed at least N times, leading the experimenter to conclude erroneously that it is representative. The probability of this is the risk α.
○ An inconclusive result, where the experimenters are not able to draw a conclusion at all. We will call this the risk β.

Note that the risks α and β are defined as analogies to the classical hypothesis testing framework. In that framework, α represents the risk of accepting a wrong conclusion (by rejecting the null hypothesis, even though it is true) and β is the risk of failing to reach a conclusion (by failing to reject the null hypothesis, even though it is false). The present framework doesn’t explicitly distinguishes null and alternative hypotheses (instead, a ‘null result’ is one outcome amongst others). However, our definitions of α (the risk of reaching a false conclusion) and β (the risk of obtaining an inconclusive result) relate to situations that are similar as in classical hypothesis testing.

### Simulations

Given a prior distribution, we can compute the risks α and β associated with any N-oo-M model by performing rather straightforward Monte Carlo simulations. We provide a simple Matlab script in Appendix.

## Results

### Example 1: two possible outcomes

We first consider a simple case where the experiment can have 2 outcomes, A and B. A is the representative outcome, with a probability of 90%, and B is an outlier with a probability of 10% (**Fig. 1A**). Because this example includes only two outcomes, it is possible to enumerate all possible results when recording from 1 to 4 animals (**Fig. 1B**): we will discuss the risks α and β of all possible models with up to 4 animals (**Fig. 1C**).

**Figure 1:**
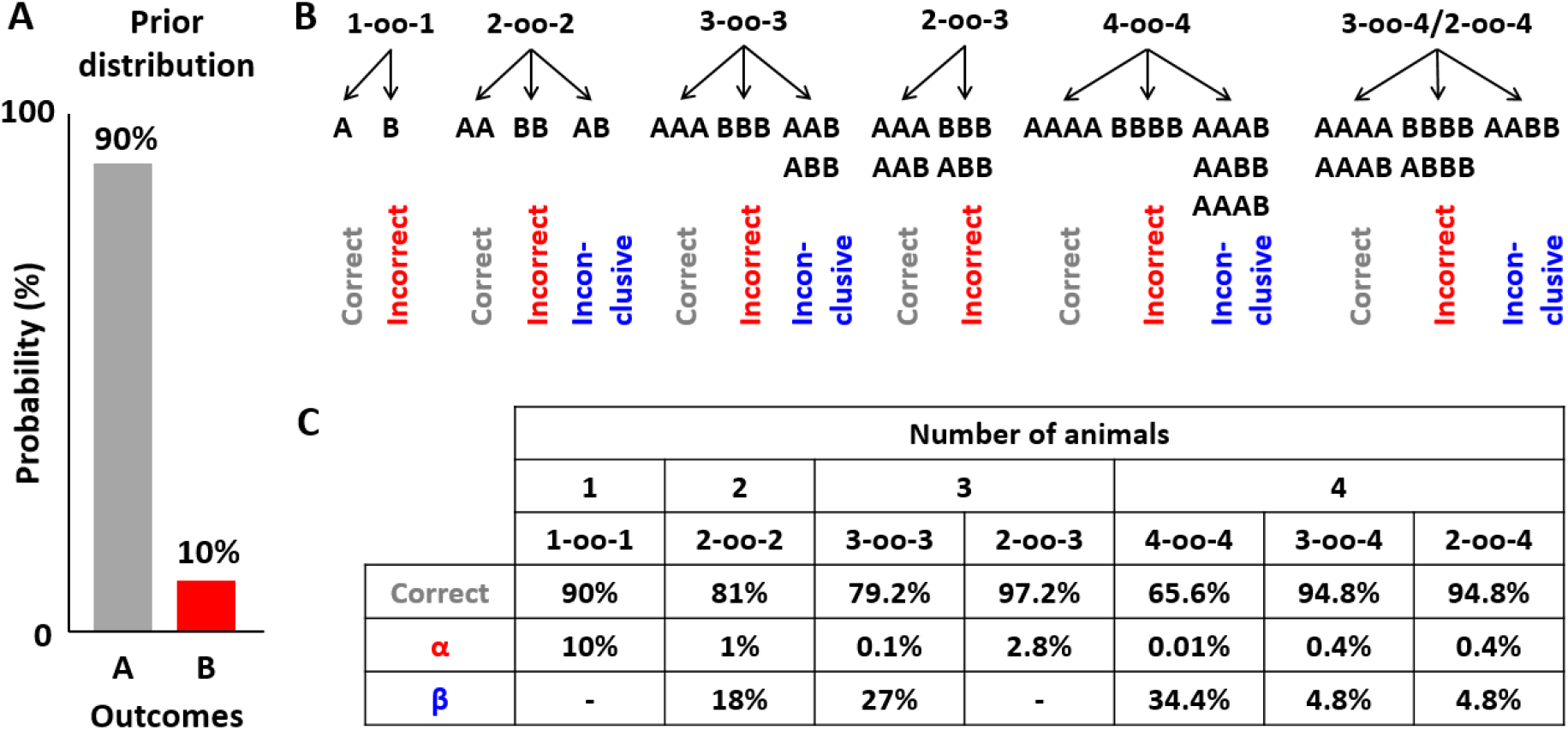
**Example 1** with two possible outcome. **A:** prior distribution. **B:** representation of all possible combinations of outcomes for all models including 1 to 4 animals. **C:** risks α and β associated with all models.

*1-oo-1 model:* with a single animal, the probabilities are evident. The experimenters have a 90% chance of observing the A, which leads to a correct conclusion, and a 10% chance of observing B, which leads an incorrect conclusion. The risk α is therefore 10%. It is impossible to reach an inconclusive result with the 1-oo-1 model and therefore β is zero.

We can conclude that the 1-oo-1 model is very problematic, since it leaves a high risk α of 10%.

*2-oo-2 model:* with two animals, there are three possibilities: AA, which leads to a correct conclusion, BB, which leads to an incorrect conclusion, and AB, which is inconclusive. The risk α of associated with BB is strongly reduced (to 1%) compared to the 1-oo-1 model, since it is equal to the probability of observing B twice, which is 10% squared. However, the probability of AA is only 81% (the square of 90%), leaving a risk β of 18%.

This indicates that the classical approach of recording from two animals reduces α drastically, but leaves substantial β risks.

*3-oo-3 model:* in this model, the experimenter will only accept a conclusion if it is supported by a unanimous outcome in three animals. This reduces the risk α to a 0.1% (i.e. the probability of BBB). However, the probability of observing AAA is only ~72% (i.e. 90% cubed), leaving an even larger risk β of 27%.

We can conclude that the 3-oo-3 model is a poor choice in most situations. By imposing a stringent criterion of 3 identical outcomes, it creates a large β risks while reducing α to a value that is probably unnecessary low. But what happens if that stringent criterion is relaxed by accepting two identical outcomes as conclusive, i.e. by using the 2-oo-3 model?

*2-oo-3 model:* when recording 3 animals, we may observe 4 possible combinations: AAA, AAB, ABB or BBB. The first two lead to a correct conclusion and the last two to an incorrect conclusion (there is no inconclusive result in this particular case). Strikingly, the risk α (of observing ABB or BBB) is only 2.8% in this example, and therefore the model is correct 97.2% of the time!

Thus, in this example, the 2-oo-3 model strikes a very interesting balance of acceptable α and low β risk. As we will see in the next section, β is not always zero but tends to remain low when using the 2-oo-3 model.

*2-oo-4 and 3-oo-4 models:* We will conclude this section by considering models using 4 animals. The 4-oo-4 model leads to an excessive β risk (compared to the 3-oo-3 model) and is not worth discussing further. In this example, the 3-oo-4 and 2-oo-4 models are identical, since AAAA or AAAB will lead to a correct result (with 94.8% probability) and BBBB or ABBB will lead to an incorrect result (with α = 0.4% probability). Finally, AABB is inconclusive with both models (in the case of the 2-oo-4 model, because it is a draw) and has a probability of β=4.8%. In this example, recording 4 animals allows decreasing α even further while maintaining a low β.

### Example 2: multiple outliers

Let us now consider a more complex and challenging example where the probability of the representative outcome A is lowered to 80%, and 2 additional outliers are added each with 5% probability (**Fig. 2A**; note that this corresponds to the example we used in ‘Statistical Model, Hypothesis 2’). Enumerating all possible combinations of outcomes (as in **Fig. 1B**) would be excessively lengthy; instead we will examine the risks α and β. Interestingly, the conclusions drawn from our previous example remain relatively un-altered.

**Figure 2:**
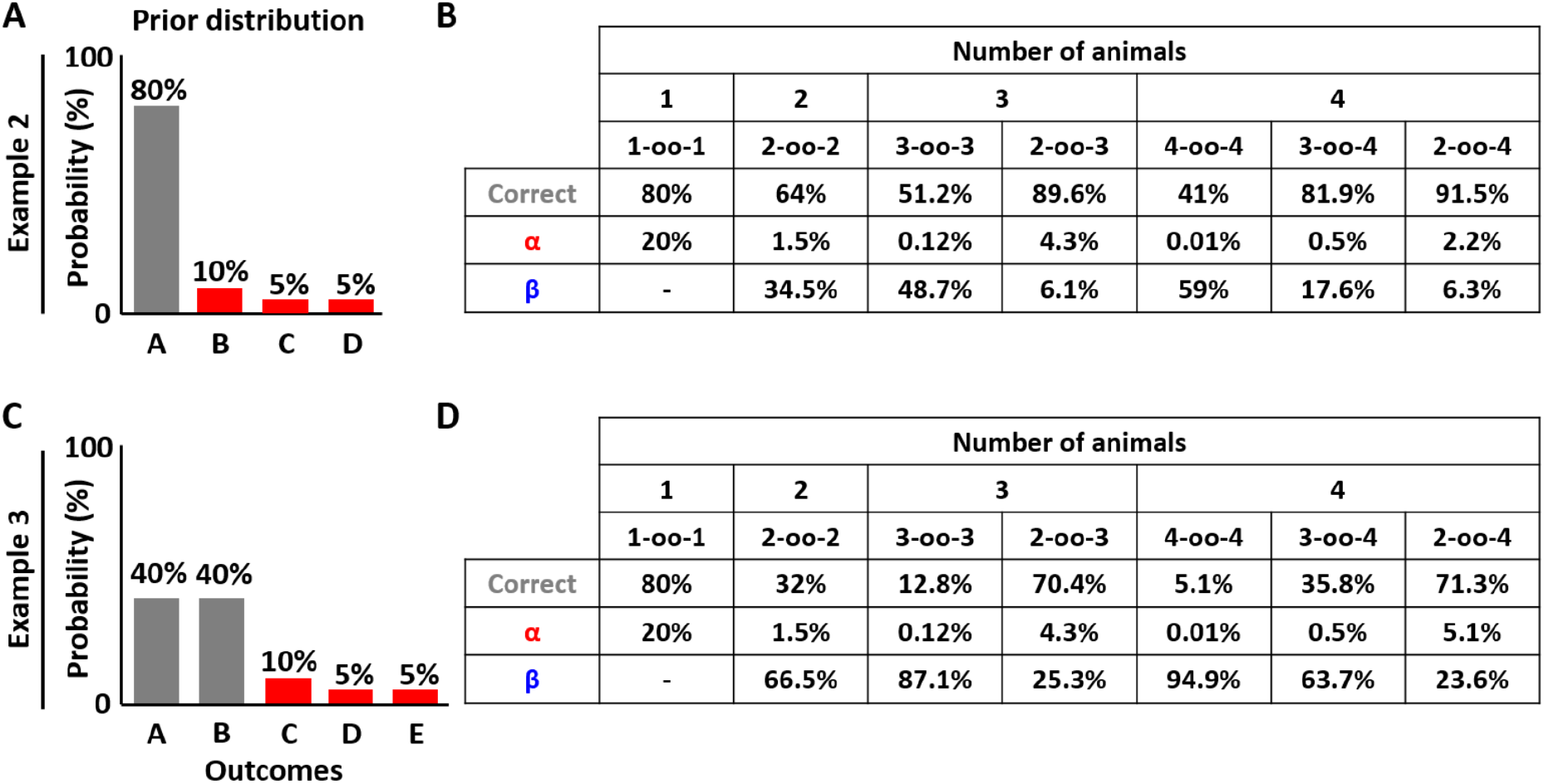
Examples 2 and 3. **A, C:** prior distributions. **B, D:** risks α and β.

*1-oo-1 model:* when multiple outliers exist (B, C and D), experimenters will reach an incorrect conclusion if they observe B, *or* C, *or* D. As pointed out in the introduction, this implies that the risk α is equal to the *cumulative* probability of all outliers, i.e. 20%. This illustrates why the 1-oo-1 model is extremely problematic when multiple sources of error exist.

*2-oo-2 model:* similar to the previous example, using two animals reduce α considerably (to 1.5%). Note that, in example 1, the risk α of the 2-oo-2 model was the square of the risk α of the 1-oo-1 model, i.e. 1% versus 10%. Here, the risk is reduced comparatively more: even though the 1-oo-1 model has a risk α of 20%, the 2-oo-2 model has a risk of only 1.5% (and not 4%, which is the square of 20%). This is because erroneous conclusions are not reached simply when recoding from two outliers, but only when recording twice from the *same* outlier. Thus, recording from two animals reduces the risk α drastically. However, β is very high (34.5%).

*3-oo-3 model:* similar to Example 1, the 3-oo-3 model reduces α even further compared to the 2-oo-2 model (to 0.1%), but also increases β (to 48.7%).

*2-oo-3 model:* similar to Example 1, we observe that the 2-oo-3 model strikes a reasonable balance: α is 4.2%, whereas β is 6.1%.

*3-oo-4 model:* this model archives a very small α (0.5%) while leaving β rather high (17.6%) but not as much as the 2-oo-2 or 3-oo-3 models.

*2-oo-4 model:* this model reduces α even further compared to 2-oo-4 (2.2%) while leaving β low (6.3%).

This example confirms the main results of Example 1, which are that recording from two animals reduces the risk α drastically, but leaves a large risk β; and that a 2-oo-3 strategy balances α and β risks.

### Example 3: multiple correct outcomes

The first two examples have assumed that only one outcome is correct. But what happens if multiple distinct outcomes are sufficiently frequent in a population that they may be considered representative? This may happen, for instance, if animals develop multiple distinct strategies to solve a task, or when animals express diverse phenotypes (e.g. eye colour).

To address this question, we need to generalize the model. With only 2 or 3 animals, it is clearly impossible to *conclusively* identify *both* outcomes. Instead, we consider that the experiment’s conclusion is correct if *one* of the representative outcomes is identified.

We simulate an experiment where the prior distribution (**Fig. 2C**) includes two representative outcomes (A and B) with a probability of 40% each, and multiple outliers (C, D, E), totalling a probability of 20%. This is similar to Example 2, except that the representative outcome has been replaced by two equiprobable outcomes.

*1-oo-1 model:* in this model, observing A or B leads to a correct result: this has a probability of 80%. Observing any of the outliers leads to an incorrect results: this has a risk α is 20%.

*2-oo-2 model:* the risk α is identical as in Example 2 (1.5%). However, it is now quite likely to observe one A and one B, which is inconclusive. As a consequence, β has a value of 66.5%, and the probability of reaching a correct conclusion is only 32%.

*3-oo-3 model:* similar to previous examples, α is very low (0.1%), but β increases to 87%.

*2-oo-3 model:* once again, the 2-oo-3 model strikes a reasonable balance: α is 4.3% (identical to example 2), and the risk β is 25.3%. While this is far from negligible, the probability of obtaining a correct answer is 70.4%.

*3-oo-4 model:* this model archives a very small α (0.5%) but still a high β (63.7%).

*2-oo-4 model:* this model performs similarly to the 2-oo-3 model: α is 5.1% and β is 23.6%. It is interesting to consider the special case of ‘ties’ in this model. Indeed, in the original formulation of the 2-oo-4 model, observing two A and two B would be considered a tie. One may alter the procedure and conclude that both A and B are representative, leading to a correct conclusion. If this procedure is followed, the risk β decreases to 5.3%. However, this increases α to 8%. While problematic in this case, this version of the 2-oo-4 model may be an option when the percentage of outliers is lower.

As a conclusion, Example 3 is particularly adversarial: one animal in 5 is expected to be an outlier, and the remaining animals are divided in two categories. And yet, the 2-oo-3 model archives a remarkable performance: with only 3 animals, it can identify one representative outcome with a probability of 70%, while keeping the risk α to an acceptable level.

## Discussion

Statistical power calculations are a crucial element of experiment planning, data analysis and interpretation. Until now, few studies had attempted to formalize the statistical power of experiments involving small numbers of animals (but see REF^1,2^, discussed below). Here, I have developed a model that applies to a variety of situations, and in particular to neuroscience studies in NHPs. This model is implemented using a short Matlab script provided in Appendix. Strikingly, the general conclusions of the model are relatively constant across a variety of scenario. They can be recapitulated as follows:

○ *One animal:* the fundamental issue with recording from only one animal is the risk of selecting an atypical animal, or performing an experimental error, leading to an incorrect conclusion. This is defined at risk α here. This may occur because individual animals exhibit abnormal anatomy, physiology or behaviour; or because of experimental factors: these sources of errors are collectively referred to as outliers in the present article. Critically, when multiple outliers exist, their probability accumulate and can potentially create high α risks. In brief, if 2% of animals have atypical anatomies, 2% aberrant behaviours, and if mistakes occur in 2% of experiments, then the total risk α is about 6%. This justifies a common rule of thumb that requires findings to be replicated in at least a second animal.
○ *Two animals:* recording from two animals reduces α dramatically, as illustrated in all the three examples (**Figs. 1–2**). Indeed, erroneous conclusions will only be reached if the experimenter records twice from *the same type* of outliers. However, when recording from two animals, there is a substantial risk of observing two different outcomes, in which case the experiment will be inconclusive (we define this as a β risk in the present framework).
○ *Three animals:* when recording from three animals, one can design a very powerful strategy by accepting results that are observed in at least two animals (‘2 out of 3’ strategy). Simulations indicate that this balances both α and β risks in a wide variety of scenario.
○ *Four animals:* interestingly, the simulation don’t demonstrate a sizeable change in statistical power when using four animals, at least within this framework. Note, however, that using four animal or more may be valuable when aiming at quantitative measurements.

In conclusion, this model supports the rule of thumb that results observed in two animals are statistically sound, but suggests that including three animals can markedly increase a study’s power.

### Two or three?

The advantage of the 2-oo-3 model is that is reduces the risk β risks (of inconclusive results) compared to the 2-oo-2 model.

In the 2-oo-2 model, inconclusive results happen when distinct outcomes are observed in two animals. On possible strategy to mitigate this risk could be to record from two animals first, and include three animals if the outcomes of the first two are different. One can easily demonstrate that the risks α and β associated with this strategy are the same as in the 2-oo-3 model, whereas the number of animals is reduced (on average, it is 2+b, where b is the β risk of the 2-oo-2 model). This is therefore a viable strategy.

The drawback of this strategy is that it will not function if one wishes to perform further analysis when the experiment is finished and the opportunity to record from a third animal no longer exists. As open-access data become the norm, providing three animals or more from the onset ensures that the necessary statistical power is available to test hypotheses that have not been formulated yet. Ultimately, the possibility to conduct further analyses at a later time reduces the total number of animal experiments, and may offset the additional experiments performed when recording from three animals. There is therefore a statistical (and ethical) justification for aiming at including 3 animals, or even more, when feasible.

This being said, it is also clear that, once a result has been observed in two animals, it may be considered statistically sound; and investigators may also chose to stop experimenting in order to save laboratory time and resources.

### Conjunction analysis framework

In REF^1^, Fries and Maris approach the question of recordings in small number of animal from the point of view of a framework named conjunction analysis^2^. This framework shows how to compute a confidence interval for the frequency of an outcome (i.e. how to infer the prior distribution), given that it has been observed N out of N time. This confidence interval takes the form of [γ_c_ 1], where 1 is the upper bound (this is trivial) and the lower bound γc follows this formula (adapted to match the notations of this article):

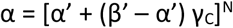

in which α’ and β’ are the experiment’s false positive and false negative rate within each subject. If we assume that α’=β’= 0 (see footnote ^1^), this simplifies to γ_c_ = α^1/N^, which is easily explained. If an outcome has a prior probability γ, then the probability of observing it N out of N times is γ^N^. If γ is unknown, we can infer with a certain risk α that γ^N^ >= α, leading to the lower bound γ_c_ =α^1/N^. For instance, if the outcome is observed in 1 out of 1 animal, we can infer that it is representative of at least γ_c_ = 5% of the population (at α=5%). If we observe it in 2 out of 2 animals, we can infer that it is representative of at least γ_c_ = 22.4% of the population. For N=3 and N=4, γ_c_ has a value of 36.8% and 47.3% respectively.

Note that simulations based on the N-oo-N model would lead to equivalent results. Therefore, from a mathematical point of view, there is no discrepancy between conjunction analysis and the present formalism.

Thus, observing the same outcome in two animals only “establishes” (with a risk α=5%) that this outcome is representative of at least 22.4% of the population. In REF^1^, Fries and Maris argue that this is not truly more useful than the lower bound of 5% obtained with 1 animal. I would diverge from this point of view for two reasons.

First, 5% represents one animal out of 20 whereas 22.4% represents more than one out of 5. If a scientist identifies a novel result, and subsequent studies show that this result holds in only in 22.4% of the population, then the initial results will, at least, be representative of a sizeable fraction of the population. In this respect, recording from 2 animals is more useful than recording from 1.

Second, it is true that we can’t infer the frequency of an outcome accurately based on 2 or 3 animals. But we can, with an acceptable degree of certainty (i.e. a low risk α), rule out aberrant results that are due to experimental errors or abnormal animals (i.e. outliers), as long as these results don’t occur in more than ~20% of the population.

Thus, the usefulness of recording more than one animal lies in reliably identifying outcomes that are present in sizeables fractions of the population, and reliably ruling out outliers.

### Choosing a prior

One limitation of the “N-oo-M” framework is that it requires assuming a prior distribution. While this is a limitation, we should keep in mind that the need of assuming a prior is inherent to any statistical power calculation performed before performing an experiment. Furthermore, we saw that the overall conclusion of the N-oo-M model are well conserved across a variety of priors. Finally, the N-oo-M framework is mathematically consistent with the conjunction analysis framework^2^, which allows inferring the prior distribution based on the results. Thus, these two approaches may complement each other.

## Conclusion

In non-human primate neuroscience, performing experiments in two or three animals has been a longstanding rule of thumb. However, formal statistical support for this practice has been lacking. The framework shown here answers this gap, and demonstrates the advantages of using two, three or more animals.

Even though using two or three animals may efficiently weed out aberrant results, the statistical power obtained with small samples remains limited. For instance, a study in two animals may identify a result that is only representative of about a fifth of the population. In cases where many outcomes are possible (for instance when measuring eye colour in humans), it is evident that obtaining a full and accurate estimation of the underlying distribution requires a large sample. These limitations must be acknowledged: the N-oo-M model can’t (and doesn’t claim to) create statistical power out of thin air. Its role is to quantify, formally, what statistical power exists in small samples.

The material and ethical challenges of neuroscience research in NHPs are considerable. There will always be circumstances where data from only one animal is available. The ethical cost of obtaining data from one animal is significant, and one can reasonably argue that there is value in publishing it, as long as it is understood that there is a larger risk α associated with such results. In the general case, there is a strong statistical argument for aiming at including two or three animals in NHP studies.

## Appendix: Matlab code

~~~
% Specify the prior distribution, and which outcomes are representative or outliers
Prior = [0.9 0.1] ; Representative = [1] ; Outliers = [2] ; % This is Example 1
Prior = [0.8 0.1 0.05 0.05] ; Representative = [1] ; Outliers = [2 3 4] ; % This is Example 2
Prior = [0.4 0.4 0.1 0.05 0.05]; Representative = [1 2] ; Outliers = [3 4 5] ; % This is Example 3
%Enter the parameters N and M of the N-oo-M model
N = 2; M = 3;
% Number of samples in the Monte-Carlo simulation
n_boot=10000000 ;
X = rand(n_boot,M);Outcome = X*0 ;
C = cumsum(Prior);
for i = 1:length(C)
    Outcome(X>C(i))=i;
end
% This lists the outcomes for n_boot repetitions and M animals
Outcome=Outcome+1 ;
n_outcomes = length(Prior) ;
S =zeros(n_boot,n_outcomes) ;
for i = 1:n_outcomes, S(:,i)=sum(Outcome==i,2);end
Inconclusive = sum(S>=N,2)~=1 ;
Correct = any(S(:,Representative)>=N,2) & ~Inconclusive ;
Incorrect = any(S(:,Outliers)>=N,2) & ~Inconclusive ;
Probability_correct = sum(Correct)/n_boot ;
Risk_Alpha = sum(Incorrect)/n_boot ;
Risk_Beta = sum(Inconclusive)/n_boot ;
~~~

## Footnotes

1 In REF^1^, Fries and Maris use α’ = 0.05 and β’ = 0, leading to γ_c_ = 18.3%, 33.5% and 44.5% for N = 2, 3 and 4 animals. Since these values are close to the one discussed here, we can conclude that our formalisms are in general agreement.

